# Females recruit faster alpha motor neurons than males during the patellar stretch reflex: evidence for neurophysiological sexual dimorphism

**DOI:** 10.1101/202713

**Authors:** Georgina Tindell, Paul A. Smith

## Abstract

**Objective:** Human neurophysiology is sexually dimorphic where the pre-motor latency of the patella stretch reflex (PR) in females occurs earlier than males; an effect unaccounted for by stature difference. Using surface electromyography we investigated whether this difference is due to dimorphism in the conduction velocity (MNCV) of the associated alpha-motor neurons.

**Methods:** Extracellular linear electrode arrays measured vastus lateralis (VL) motor unit action potential activity (MUAP). MNCV was determined from the relationship between the latency of MUAP initiation and VL distance during a PR.

**Results:** In females, the PR premotor latency occurred 2.1±0.3 ms earlier than in males and recruited alpha-motor neurons with a MNCV 5 ms^−1^ faster (p<0.02) than males: 52.4±1.5 5 m s^−1^ (n=40) compared to 47.6±1.3 5 m s^−1^ (n=41) respectively.

**Conclusions:** In males the longer PR premotor latency was accounted for by their slower MNCV. Our finding is contrary to the observations of larger diameter alpha-motor axons and somas seen in males. Since the PR yields MNCVs values at the lower end of published values supports the “size-principle” of motor unit recruitment with a physiological stimulus.

**Significance:** Our findings evidence gender dimorphism in a routine neurophysiologic test which suggest androgen modulation of skeletal muscle innervation.

## 1. Introduction

In humans the patella stretch reflex (PR) or “knee-jerk reflex” is widely used to aid the diagnosis of neurological dysfunction. On stretch stimulation of the muscle spindles within the quadricep muscle group, electrical information is relayed via 1a afferent sensory nerves to the L3-L4 region of the spine. From here, efferent motor output is elicited in the α-motor neurons that innervate the quadricep muscles which causes them to contract and extend the lower leg (Schubert and Kiel, 1968). Physiologically, the PR forms part of a series of circuits that underlies the tone and function of the locomotor system. Electrophysiologically, the PR can be quantified in terms of the associated motor unit action potential (MUAP) activity. Interestingly in human females the temporal delay from the stretch stimulus to the appearance of MAPs, the pre-motor latency, occurs earlier than in males; an observation unaccounted for by differences in stature between genders (Vickery and Smith, 2012) but one that may relate to other neuroanatomical differences.

Sexual dimorphism of human neuroanatomy is well established (Morris et al., 2004). Although such differences primarily relate to the biology of reproduction, such as the motor neurons of the spinal nucleolus of bulbocavernosis which are of larger size and number in males than for females (Forger and Breedlove, 1986), other differences do exist. For example, the cross sectional areas of the median and sciatic nerves are larger in males than females (Ide, 1930). The neurons involved in the PR are no exception to this, for example, the soma of participant motor neurons in the ventral horn of L3-L4 are larger in males than females (Yuan et al., 2000); a structural characteristic canonically expected to yield faster nerve conduction velocities for males, an idea that apparently contradicts the finding that the PR pre-motor latency is longer in males than females (Vickery and Smith, 2012). The mechanical aspects of the PR also display clear gender differences; phenomena that closely relate to differences in the distribution and size of muscle fibre types of the quadriceps between genders (Sadoyama et al., 1988; Viitasalo et al., 1981; Staron et al., 2000; Simoneau and Bouchard, 1989).

Based on biophysical considerations (Kernell, 1966; Lüscher et al., 1979; Smith and Vickery, 2012) a canonical explanation for the longer PR premotor latency in males is the “size principle” of recruitment (Mendell, 2005). Since larger motor neurons require greater amounts of synaptic input and summation to elicit excitation (Clamann and Henneman, 1976), this may be delayed in males. Indeed, females show a more varied motor unit recruitment during low intensity exercise thought indicative of gender differences in spinal integration (Nishikawa et al 2017). However, an alternative explanation is that a unique population of α-motor neurons are recruited during the PR whose MNCV is less related to somal size and axon diameter but more dependent on other factors such as internodal distance, myelination, or ion channel density (Waxman, 1980). Whether gender differences in the MNCV of motor nerves recruited during the PR accounts for their pre-motor latency difference is unknown. Although the mean femoral motor nerve conduction velocity has been measured to range between 60-70 m/s (Shubert and Kiel, 1968; Chopra and Hurwitz, 1968; Gassel, 1963; Uludag et al., 2000) these were from mixed gender samples. Of interest is that the MNCV with the median and ulnar nerve trunks are positively correlated with oestrogen level (Singh et al., 2016) given the marked difference in the level of this androgen between genders one can predict faster MNCVs in females. Indeed, no study that specifically addresses gender effects on human neurophysiology has been published to date, an especially pertinent fact given the current interest in this topic (Cahill and Aswad, 2015).

The aim of this study was to determine if either the conduction velocity of the motor nerves (MNCV) recruited during the PR demonstrates gender dimorphism in humans and if so whether these are sufficiently large enough to account for the gender difference in pre-motor latency. In this report, we demonstrate that the motor nerves responsible for VL muscle excitation during the PR are substantially slower than that reported for extracellular stimulation studies of the femoral nerve, and ~5 m/s faster in females than in males; a difference unrelated to stature and one that can entirely explain the difference in premotor latency between the two sexes. Moreover, the methodology employed exonerates any spinal contribution to the gender difference in PR latencies.

## 2. Methods

### 2.1 Experimental rationale

Previous measurements α-motor MNCV of the femoral nerve used extracellular stimulation of the nerve trunk at the point where it emerged from the inguinal ligament (c.f. Schubert and Kiel, 1968; Gassel, 1963). Electromyography then detected motor nerve conduction via the appearance of MUAPs from the respective innervated quadriceps muscle with electromyography; a technique that takes advantage of the high nerve/muscle innervation ratio of ~ 1500 (Rich et al., 1998) to improve the signal to noise ratio. The MNCV is then calculated from the difference in pre-motor time delay, δt, between these two locations of known geometrical separation, δd, (Figure 1) by δd/δt (Shubert and Kiel, 1968; Gassel, 1963; Uludag et al., 2000). However, since neurons with the lowest threshold on extracellular electrical stimulation are those with the largest axonal diameters and fastest conduction velocities these are primarily recruited first and the MNCV is overestimated. Secondly, the alpha-motor neurons sampled are not necessarily those recruited during a PR, where motor neurons with the smaller soma and axon diameters which, according to the “size principle”, will be preferentially excited first (Lüscher et al., 1979; Mendell, 2005). Consequently, the use of a physiological stimulus via a tendon tap is preferential to external electrical stimulation in order to maintain the physiological relevance of MNCV measurement.

**Figure 1.**
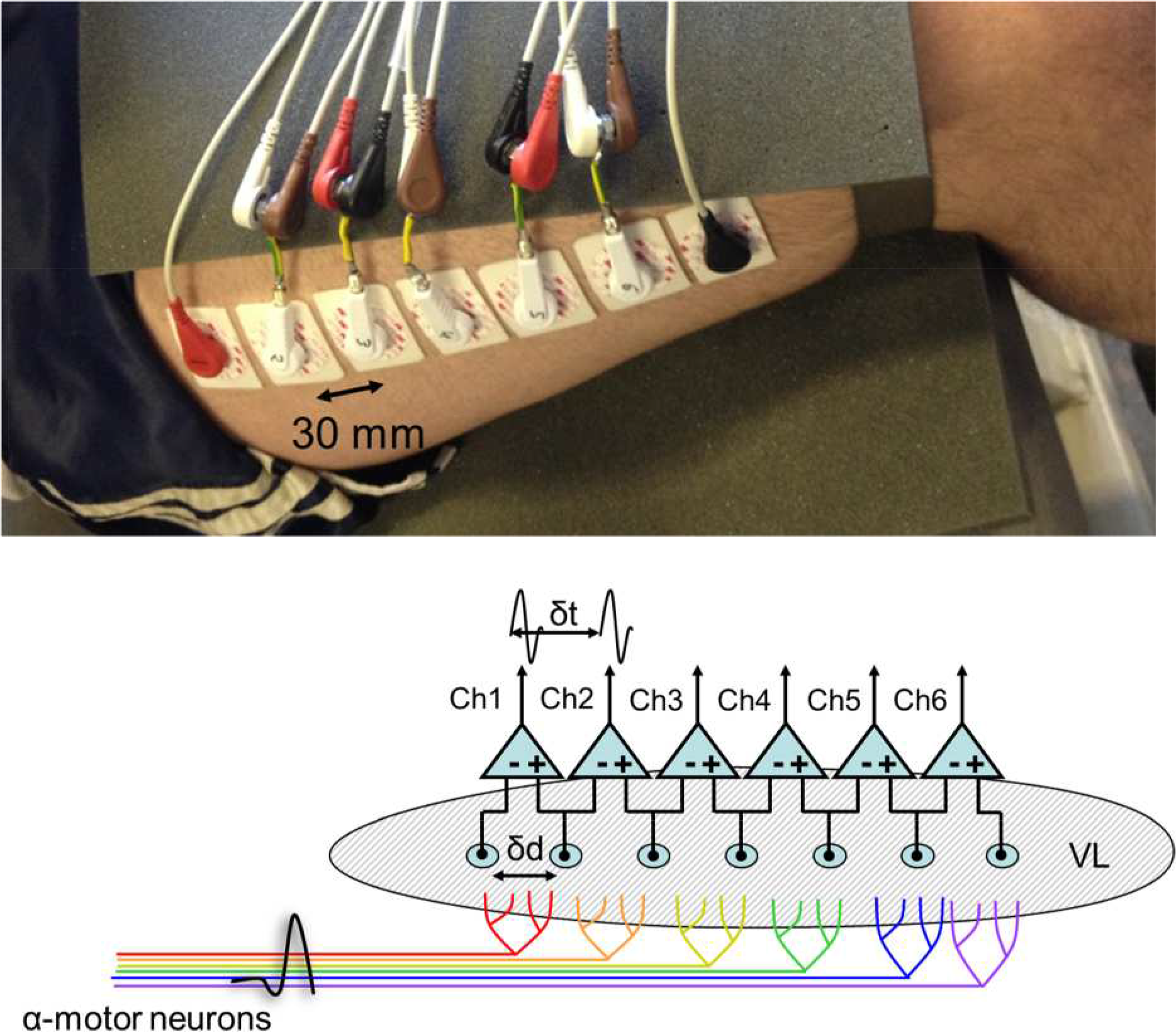
Arrangement of electrodes for sEMG measurement from the vastus lateralis (VL). a) Image of a subject’s right leg which illustrates the placement of seven electrodes as well as the earth strap above the knee. Note that the five central electrodes have split outputs which feed into shared inputs of the differential amplifiers of the acquisition device. b) Schematic of the flow of electrical information during a PR. The inter electrode separation is given by δd, whereas the latency with proximal conduction is given by δt..

To calculate α-motor MNCV during the PR, values of δd and δt are required. To do this with a physiological stimulus, we took advantage of the fact that the innervation zones (IZ) of the VL are not just clustered at the muscle belly of the VL (Masuda and Sadoyama, 1987) but are dispersed along its length (Saitou et al., 2000), as well the fact that the distal branch of the femoral nerve that serves the VL runs proximally to distally for the majority of its length (Patil et al., 2007). Consequently during the time course of a PR it is assumed a wave of excitation down the α-motor neurons of the VL is mirrored by a wave of muscle innervation and excitation of MUAPs down the VL. The appearance of MUAPs was detected by a linear array of surface electrodes. The relationship of the time delay, δt, between the appearance of MUAPs at successive electrodes and their known geometric spacing δd allows estimation of the underlying MNCV of excitation.

### 2.2 Subjects

Subjects were näive to the experimental method. In total 81 subjects enrolled in the study, 41 male and 40 female with an age range between 19 and 21 years old; anthropometric details in Table 1. After explanation of the protocol, each subject read a human volunteer information sheet and answered at medical questionnaire; subjects were excluded if they had any history of neurological or neuromuscular disease or lower limb injury. The purpose of the study was explained and informed written consent was taken, subjects were free to withdraw at any point. The study conformed to the codeset by the Declaration of Helsinki and was approved by the ethical committee of the Faculty of Medicine, University of Nottingham.

**Table 1.** Anthropometric data of the study subjects, data is given as means ± S.E.M. (minimum - maximum).

| **Gender** | **Height (m)** | **Thigh Length (cm)** |
| --- | --- | --- |
| **Male (n = 41)** | 1.79 ± 0.95 (1.69-1.91) | 51 ± 0.59 (43-59.5) |
| **Female (n = 40)** | 1.67 ± 0.82 (1.54-1.78) | 47.4 ± 0.5 (42-55.5) |
| **Mean Difference** | 0.12 ± 0.01 | 3.55 ± 0.78 |
| **P (Unpaired t test)** | P<0.0001 | P<0.0001 |

### 2.3 Experimental protocol

Subjects were seated on an electrically insulating foam cushion with their upper legs horizontal and their lower leg hung at ~110° flexion, as determined with a protractor. Electrode pads (Red Dot, 3M) were placed along the longitudinal axis of the VL muscle on the dominant leg (determined by preference in sport). Prior to electrode placement, the skin was wiped with 70% Isopropyl alcohol and abraded to reduce electrical impedance. Two arrangements of electrodes were used; in the first, an electrode was placed at 5/6 of the measured difference from the anterior superior iliac spine (ASIS) to the lateral side of the patellar. A further 6 electrodes were then placed proximally with a 30 mm inter-electrode distance (IED) to create a linear array of 7 electrodes along the VL (Fig. 2a). In the second arrangement, a pair of electrodes were placed to straddle 2/3 of the measured difference from the ASIS to the lateral side of the patellar with an additional electrode placed proximally and two more distally, but all with a 40 mm IED, hence creating a linear array of 5 electrodes. A reference strap electrode was attached around the ankle and calf. These arrangements form an electrode array similar to those previously decribed (Rainoldi et al., 2000; Saitou et al., 2000) but with a largerer IED seperation as to maximize the inter electrode sample time and reduce the degree of crosstalk via secondary sampling of fascicles served by neighouring electrodes.

**Figure 2.**
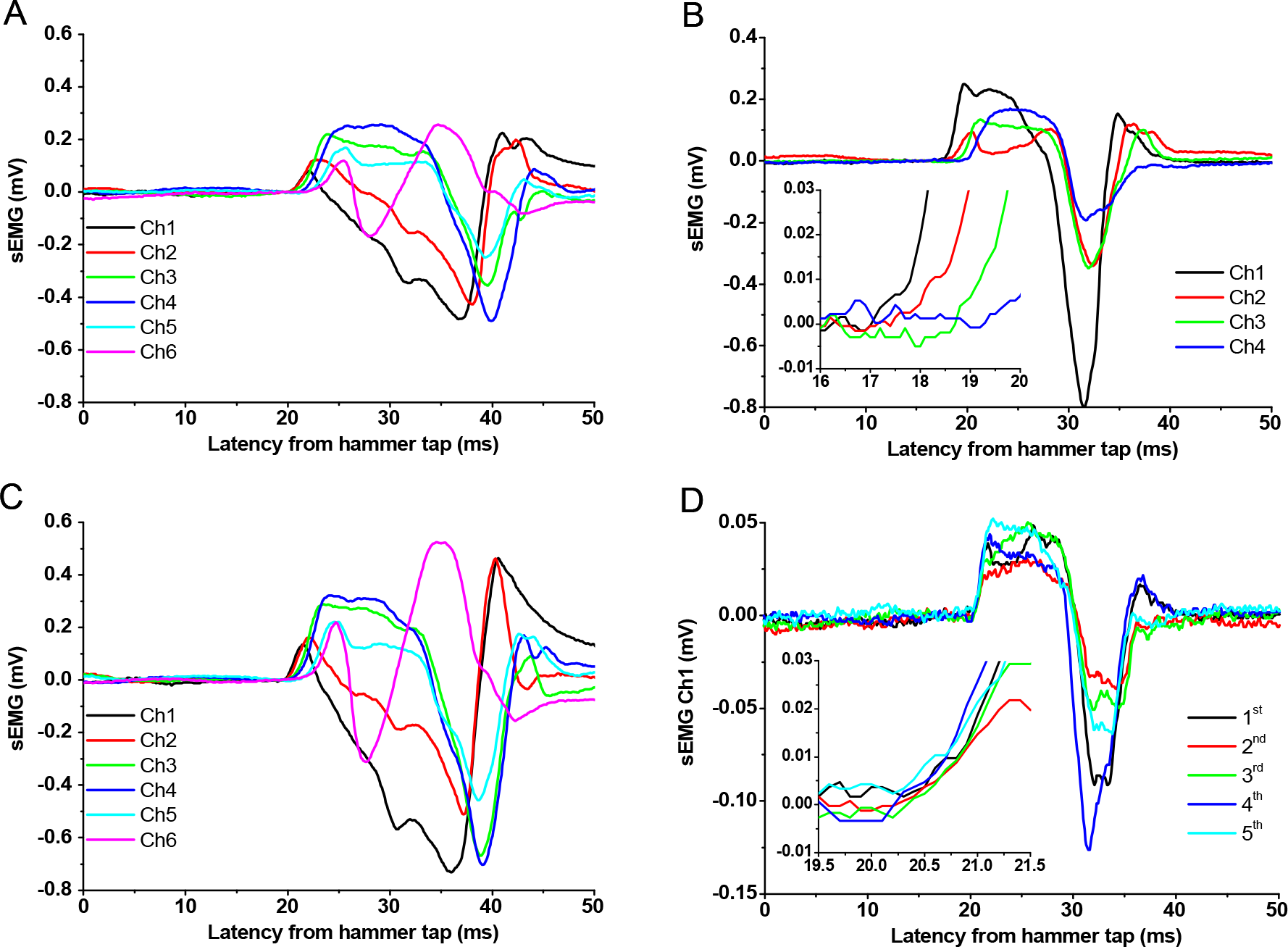
Representative sEMGs recorded from the vastus lateralis during a patellar reflex elicited at time zero for two different female subjects. A and C) sEMGs recorded from a 7-electrode array with an inter electrode distance of 30 mm in response to a hammer tap of 125 mJ (A) and then 450 mJ (C). Traces Ch1-Ch6 are recorded proximally to distal respectively. Note the similarity in the shape of the sEMG waveform for any given electrode pair when elicited at the higher energy. B) sEMGs recorded from a five electrode array with an inter electrode distance of 40 mm in response to the hammer tap of 200 mJ. Traces Ch1-Ch4 are recorded proximal to distal respectively and reveal a clear temporal separation of MUAP transit. D) sEMGs recorded from electrode pair Ch4 in the same subject as in B) but in response to 5 consecutive PRs labelled 1 to 5. Note the variability in amplitude, but not latency, between each PR.

A handheld hammer was used to locate the most sensitive region on the patellar tendon that elicited a PR and the patellometer placed appropriately to that region. To facilitate the reflex, and to limit central modulation, the Jendrassik manoeuvre (Delwaide and Tolulouse, 1981) was applied prior to each PR: subjects were instructed to interlock their fingers and pull isometrically when asked. Hammer taps were only applied when the leg was stationary and relaxed as defined by the absence of sEMG activity. A minimum rest period of 5 s between taps limited confounders that may arise through subject anticipation and fatigue. The protocol was terminated if the subject experienced pain or discomfort. The PR was elicted by kinetic energies as calculated with the following equation:

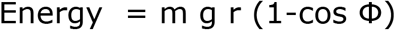

Where m is the mass of the hammer (0.178 kg), r the radius of the hammer motion (0.3 m), g the acceleration due to gravity (9.8 m s^−2^) and Φ the vertical angle from which the hammer was released. The hammer contained a piezo accelerometer whose ouput was used to trigger data aquisition on collision with the patellar tendon. The PR was eilcited by at consecutive tap to yield at least 5 estimates of latency for that subject that were readilly measurable and not masked by electrical noise or movement artefacts.

### 2.4 Data Aquisition and Analysis

Data was collected with 3 × ADI Powerlab 26Ts, which conformed to ISO9001 for human use, and aquired by Lab Chart Version 7 software (ADI Instruments, Oxfordshire, UK). To determine the presence of background EMG activity, data was collected 50 ms prior to trigger. The sEMG signal was filtered (−3 db) with a 0.5 Hz 1^st^ order high pass and a 4^th^ order 2 kHz low pass Bessel, prior to digitization at 10 kHz per channel to give a 10-90% signal rise time less than 0.2 ms.

The MUAP data was analysed for latency; determined as the time point at which the sEMG deviated postively and continuosly from the baseline as the defined upstroke of the leading edge of the MUAP (c.f. fig 2c.). For and given subject the reported latency values at any given electrode is the mean of at least 5 determinations.

Statistical analysis was performed with Graphpad PRISM version 6 (San Diego, California USA) with tests as given in the text. Data sets were tested for normality with the D’Agostino & Pearson omnibus test. Data are represented as either means ± S.E.M. or as medians with 5 to 95% confidence intervals (C.I.), with n the number of determinations. The fitting of data with equations given in the text used the least squares algorithm as supplied with PRISM. Statistical significance is defined as P <0.05 and is flagged as * in graphics, ** when P <0.01 or *** when P <0.001.

## 3. Results

Figure 2 shows representative sEMGs recorded with a 7-electrode and a 5-electrode array for two different female subjects. In both cases the MUAP latency increased with electrode distance (Fig. 2A, B & 3B). MUAP waveform profiles were variable both between subjects and electrodes, sometimes biphasic but more often polyphasic in nature. MUAPs often had waveforms profiles that appeared to share components, attenuated or/and temporally shifted, from neighbouring electrodes (e.g. compare Ch1 with Ch2, and Ch4 with Ch5 in Fig. 2A & 2C). Although successive MUAPs recorded from a given electrode pair in response to the same energy varied in magnitude, this phenomenon was neither associated with a change in waveform profile or latency (Fig. 2D inset & 3A).

For all subjects, MUAP latency was positively correlated (p<0.05 Pearson) with electrode distance and readily fitted by linear regression (solid line, Fig. 3B), where the MNCV for the underlying α-motor neuron activity is given by the reciprocal of the slope parameter (Gassel, 1963). At higher hammer energies, the MUAP increased in amplitude but with little change in waveform profile (Fig. 2C). Although this was associated with a small but significant decrease in latency (−0.55 ± 13 ms, p = 0.003, Paired t test, n =28, Fig. 3B) a similar shift was seen at all electrodes with the sequel that MNCV was unchanged (p = 0.984, Paired t-test).

**Figure 3.**
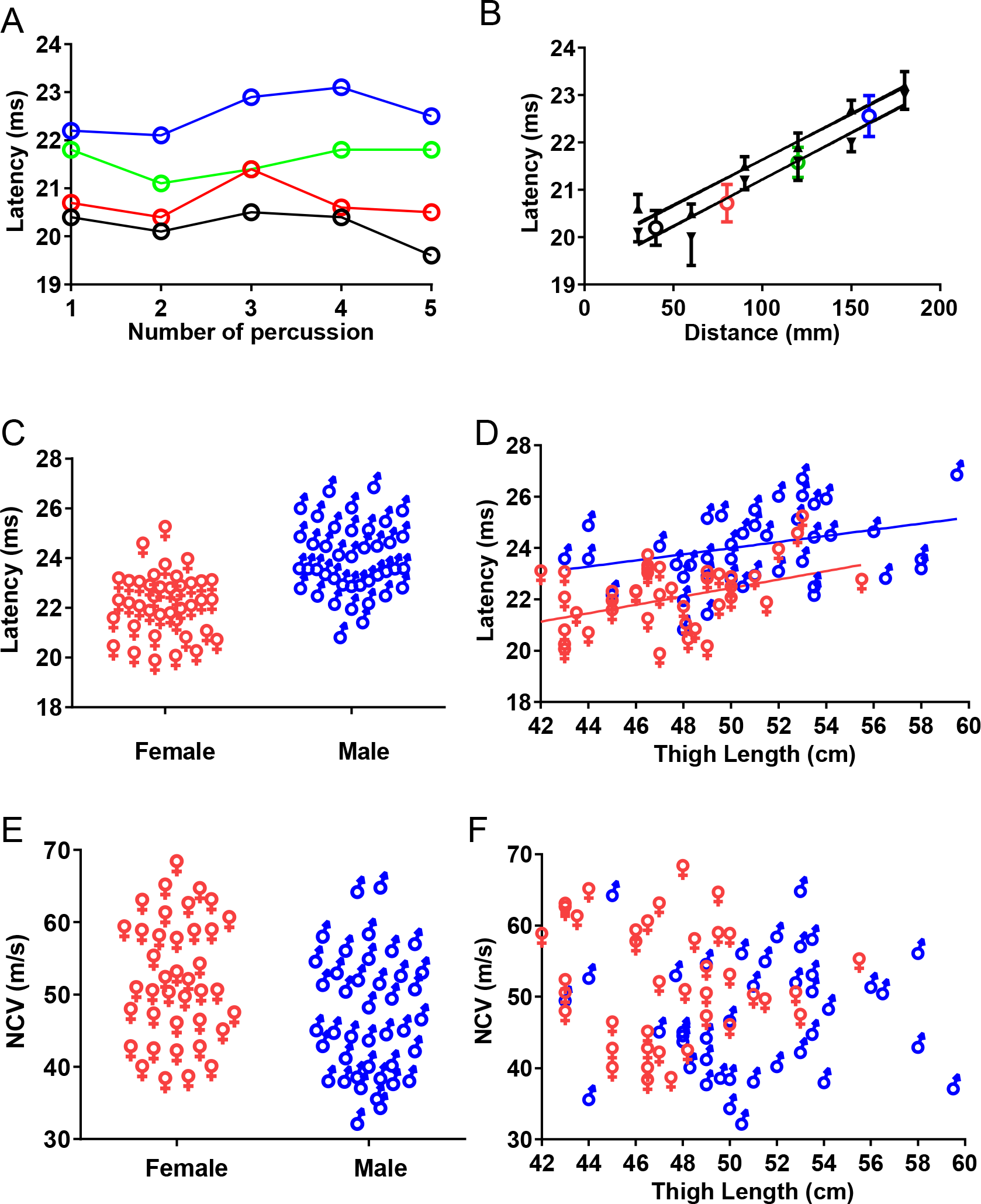
Latency measurements and estimates of motor nerve conduction velocity during the patellar reflex. A) Representative plots of latency with time for the 4 different electrode positions of 40 mm IED in a female (from data shown in Fig 2B, electrodes positioned at 40, 80, 120, 160 mm from bottom to top respectively) for 5 consecutive repeats of the PR elicited by 200 mJ. B) Representative plots of mean latency ± S.E.M. (n=5) against electrode distance for the PR elicited by: 200 mJ for the female data as in A) with 40 mm IED (O); and by 125 mJ (▾) and 450 mJ (▴) for a female subject with 30 mm IED (from data as shown in Fig 2A, C); solid lines are fit by linear regression with identical reciprocal slopes of 53 m/s. C) Comparison of PR premotor latency measured at a comparable topographical position (2/3rds the thigh length from ASIS) for 40 male (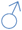) and 41 female (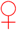) subjects (P<0.0001, t-test). D) Correlation of PR latency plotted against thigh length for 40 male (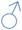) and 41 female (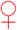) subjects. Solid lines are fit by linear regression with parameters as indicated in text. E) Comparison of motor nerve conduction velocity (MNCV) for 40 male (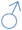) and 41 female (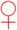) subjects (p<0.018, t - test). F) Motor Nerve conduction velocity (MNCV) plotted against thigh length for 40 male (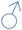) and 41 female (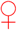) subjects. Note the lack of correlation.

The latencies measured at comparable topographical position: 2/3rds the thigh length from ASIS, for each subject is shown in Fig. 3C; occurred 2.1 ± 0.3 ms (P<0.0001, t-test) earlier for females (22.0 ± 0.2 ms, n = 40) than for males (24.1 ± 0.22 ms, n = 41).

Figure 3d shows that latency was positively correlated with thigh length for both genders; for females r = 0.42 (95% C.I. 0.127 to 0.648, Pearson P=0.0067), for males r = 0.32 (95% C.I. 0.1 to 0.57, Pearson P=0.044). These relationships were well described by linear regression with slopes of 0.16 ± 0.06 ms cm^−1^ and 0.12 ± 0.06 ms cm^−1^ for females and males respectively; values insignificantly different (P = 0.59). However, the y-intercepts were significantly different (P<0.0001, F-test) at 14.2 ± 2.7 ms and 18.0 ± 2.9 ms for females and males respectively. Figure 3D also shows that for any given thigh length males generally have the longer, slower, latency.

Figure 3E demonstrates that females had a mean MNCV of 52.4 ± 1.5 m s^−1^ (n = 40), some 5 m s^−1^faster (p<0.018, unpaired t -test) than that of males at 47.6 ± 1.3 m s^−1^ (n = 41). MNCV was not correlated with thigh length (r = 0.039, p = 0.8 and r = 0.066, p = 0.68 for females and males respectively, Pearson; Fig. 3F).

If we assume that the difference in PR latency between the genders, Δt, is primarily due to the difference in conduction time for action potential propagation down the α-motor neuron efferents then Δt can be calculated by: Δt = d_m_ / MNCV_m_ - d_f_ / MNCV_f_

Where d_m_ and d_f_ are the respective afferent path lengths from L3-L4 of the dorsal horn in males and females respectively, which we approximated to the thigh length; MNCV_m_ and MNCV_f_ are the respective MNCV for males and females, as estimated in the present study. Using this equation, Δt was calculated to be 1.69 ± 0.42 ms, a value insignificantly different to the gender difference in latency measured here (2.1 ± 0.3 ms; P = 0.34, unpaired t-test). In an attempt remove stature as a confounder these calculations were repeated using only subjects within a restricted range of 48-50 cm thigh length. This exercise yielded similar outcomes: a MNCV 8.6 ± 2.9 m s^−1^ faster for females (p=0.0071, unpaired t-test) and a Δt of 1.91 ± 0.66 ms, a value also insignificantly different to that measured (1.68 ± 0.44 ms; P = 0.74, unpaired t-test). For both scenarios the difference in MNCV between the genders readily accounts for that in latency.

## 4. Discussion

We used surface electrode arrays to measure the MNCV of the alpha-motor neurons involved during a patellar “knee jerk” stretch reflex. We confirmed that the pre-motor latency for the PR in females is significantly shorter than for males (Vickery and Smith, 2012); a phenomenon independent of stature. Importantly, we demonstrate that the MNCV responsible for the initiation of VL muscle excitation during the PR is ~5 m/s faster in females than in males; a difference that entirely explains the difference in premotor latency between the two sexes. Moreover, the methodology employed exonerates any spinal contribution to the gender difference in PR latencies. These findings are contrary to that expected from the anatomical observation that human males have larger diameter α-motor neurons compared to females (Yuan et al., 2000)

### Variation of latency with proximal distance

The similarity between inter-electrode waveforms of MUAP within a given individual is most likely due to the fields of detection of successive electrode pairs overlapping a common muscle fascicle. The muscle fascicles of the VL are 80-90mm length and arranged in a chevron, bipennate pattern with a pennation angle of 13-27° (Abe et al., 1998; Ward et al., 2009). We estimate that the maximum IED to duplicate (overlap) detection of the MUAP within a given fascicle along a longitudinal axis is somewhere less than 39 mm. This may explain why crosstalk between electrodes was more frequently observed with an IED of 30 mm rather than 40 mm.

### Conduction velocity

Our gender pooled value of 50 ± 1 m s^−1^ (n = 81) for the MNCV involved in the PR is substantially less than that measured by direct electrical stimulation of the femoral nerve: 60- 74 m s^−1^ (Schubert and Kiel, 1968; Gassel, 1963; Uludag et al., 2000; Chopra and Hurwitz, 1968). This apparent disparity in MNCV value most likely arises from the different method of stimulation we employed and the motor pool recruited as discussed below.

### Gender difference in MNCV

This is the first time a gender difference in the MNCV of α-motor nerves recruited during a physiological response has been described. However the MNCV of peroneal (52 vs 49 m s^−1^ Gregersen, 1967; 50 vs 48 m s^−1^ Pickett, 1982) and ulnar (57 vs 55 m s^−1^ Pickett, 1982; 60 vs 56 m s^−1^ Robinson et al., 1993) are all also reported to be faster in females than males. Interestingly, the magnitude of these differences, 2-4 m s^−1^, are similar to that we now report during femoral motor nerve function (5 m s^−1^).

It is well understood that MNCV is proportional to axon diameter (Goldman and Albus, 1968), however, since alpha-motor neurons at the level of L3 in the anterior horn of the spinal cord have a larger cross sectional area in males (Yuan et al., 2000) this is an unlikely explanation for the gender difference we describe for MNCV; larger somas being equated with larger axon diameters and conduction velocity (Clamann and Henneman, 1976). Other, alternative mechanisms may explain our gender difference in MNCV, for example nerves with less myelination exhibit slower conduction velocities (Waxman 1980), and so it is possible that human females have an increased myelin/axon diameter ratio. Although specific to the central nervous system, gender differences in myelin turnover are implicated in the sexual dimorphism of multiple sclerosis progression (Cerghet et al., 2006). Moreover, females may possess a larger internodal separation of the nodes of Ranvier, a factor also established to increase MNCV (Wu et al., 2012). Alternatively, females may possess voltage-gated Na^+^ channels at higher densities or/and with different gating properties than their male counterparts at their nodes of Ranvier, again a factor that is associated with increased conduction velocity as well neuron excitability (Lin et al., 2002). Since the median and ulnar MNCV are positively correlated with oestrogen level in postmenopausal women (Singh et al., 2016) the extent to which neuroanatomical gender dimorphism in the femoral motor nerves are mediated by differences in circulating sex hormones awaits further investigation.

The MNCV values we report is lower than the mean 60-74 m s^−1^, and are at the lower end of the range, measured by electrical stimulation in mixed gender sample populations (50 to 79 m s^−1^ Schubert and Kiel, 1968; 56 to 94 m s^−1^ Chopra and Hurwitz, 1968; 42 to 88 m s^−1^, Uludag et al., 2000), evidence strongly supportive of the size-principle of alpha-motor neuron recruitment during physiological excitation: the smallest, slower motor neurons being recruited first (Lüscher et al., 1979; Mendell, 2005).

### Limitations

The fact that our estimates for the MNCV for alpha-motor neurons of the femoral nerve are substantially smaller than those measured by direct electrical stimulation, might be misconstrued as an artefact through muscle conduction contribution. Such an idea can be dismissed since a delay through muscle conduction would, on average, be expected to be experienced by all the electrodes in the array and would simply cause a simple upward shift towards longer latencies for the relationship between latency and electrode position with no effect on slope or our MNCV estimates.

Conduction along the muscle fibre may, however, help explain the shape of the MUAP. The initial rising phase of the MUAP is due to the detection of the earliest action potentials (MAPs) that arise from IZs in the immediate vicinity of the electrode, whereas MUAPs that originate from more distance IZs will have a component of latency due to muscle fibre conduction.

MNCV is well established to be temperature dependent with a ~ 5% increase per degree C (Kiernan et al., 2001). Since the core body temperature of females is almost identical to that of males (Sund-Levander et al., 2002), temperature is unlikely to account for the 10% difference in MNCV, which would require a core body temperature difference of at least 2°C. Difference in skin temperature is also unlikely to account for the differences in MNCV as the femoral nerve is located deep within the thigh, furthermore gender differences in MNCV reported for other motor nerves, similar to that reported here, are apparently independent of skin temperature (Robinson et al., 1993; Soudmand et al., 1982).

### Conclusions and future perspective

In this study we provide clear evidence that alpha-motor neurons involved in the PR have a faster MNCV in females than males. This finding is contrary to the observations that motor nerve trunks of males have larger dimensions of alpha-motor nerves. However, our data does support the size-principle of alpha-motor neuron recruitment during physiological excitation with MNCVs values at the lowest end of the range determined by direct electrical stimulation. Our non-invasive protocol may provide a simple neurophysiological test to complement clinical and molecular cytogenetic tests in the study of androgen modulation of MNCV (Singh et al 2016) and muscle innervation (Forger and Breedlove, 1986). This may be pertinent in conditions such as androgen insensitivity syndrome (AIS) in which feminization of males could lead to a female MNCV phenotype through anatomical effects on motor neurons (Morris et al., 2004), conversely, a male MNCV phenotype might be observed in women who have excess androgens that arose in utero with Congenital Adrenal Hyperplasia (CAH) (Collaer et al., 2009); such observations await investigation.

## Conflict of Interest

We declare that no actual, or potential, conflict of interest in relation to the is article exists.

## Authors’ contributions

GT acquired and analysed data, and also participated in experimental design. PAS conceived the study, statistical analyses and wrote the manuscript.

